# Aging Reduces EEG Markers of Recognition Despite Intact Performance: Implications for Forensic Memory Detection

**DOI:** 10.1101/2020.10.20.339523

**Authors:** Robin Hellerstedt, Arianna Moccia, Chloe M. Brunskill, Howard Bowman, Zara M. Bergström

## Abstract

ERP-based forensic memory detection is based on the logic that guilty suspects will hold incriminating knowledge about crimes they have committed, and therefore should show parietal ERP positivities related to recognition when presented with reminders of their crimes. We predicted that such forensic memory detection might however be inaccurate in older adults, because of changes to recognition-related brain activity that occurs with aging. We measured both ERPs and EEG oscillations associated with episodic old/new recognition and forensic memory detection in 30 younger (age < 30) and 30 older (age > 65) adults. EEG oscillations were included as a complementary measure which is less sensitive to temporal variability and component overlap than ERPs. In line with predictions, recognition-related parietal ERP positivities were significantly reduced in the older compared to younger group in both tasks, despite highly similar behavioural performance. We also observed ageing-related reductions in oscillatory markers of recognition in the forensic memory detection test, while the oscillatory effects associated with episodic recognition were similar across age groups. This pattern of results suggests that while both forensic memory detection and episodic recognition are accompanied by ageing-induced reductions in parietal ERP positivities, these reductions may be caused by non-overlapping mechanisms across the two tasks. Our findings suggest that EEG-based forensic memory detection tests are invalid in older populations, limiting their practical applications.

## Introduction

Recognition is associated with positive ERP peaks across the parietal scalp around half a second to a second after a person encounters a reminder. Such parietal ERP positivities have been studied extensively in theoretical investigations of the neurocognitive processes that underlie episodic memory retrieval (Rugg & Curran, 2007; Vilberg & Rugg, 2009; Wilding, 2000), and in applied forensic settings as a tool for detection of concealed memories (Bowman et al., 2013; Meijer et al., 2014; Rosenfeld, 2020). However, these different fields have largely progressed separately, and forensic applications have not incorporated findings from theoretical work about how memory-related brain processes change in older age (Friedman, 2013; Morcom, 2016). This oversight is problematic since forensic memory detection is based on the assumption that memory-related brain activity is consistent across individuals. Here, we investigated if ERP-based forensic memory detection is affected by aging, and how such detection relates to ERPs during episodic recognition. We also assessed whether EEG oscillations that can be used as markers of recognition (Hanslmayr et al., 2016; Herweg et al., 2020; Hsieh & Ranganath, 2014; Nyhus & Curran, 2010) were influenced by ageing in a parallel way to ERPs, across both episodic recognition and forensic memory detection tests.

Forensic memory detection uses non-verbal markers of memory to determine whether a suspect knows information about a crime that would be unknown to an innocent person (Lykken, 1959). The ERP-based Concealed Information Test (CIT) involves measuring if recognition-related parietal positivities (“P300s”) are enhanced around 0.5-1s after reminders of a crime (“probes”) have been presented. CITs are often conducted to detect memories of a particular event by using central details from the event as probes, such as detecting if suspects recognise the word “ring” if they had previously stolen a ring (Rosenfeld, 2020). Crime-relevant probes are repeatedly but infrequently presented, interspersed with repeatedly presented frequent irrelevant stimuli in an “oddball” design, because subjectively rare probes enhance P300 amplitudes and facilitate memory detection (Bergström et al., 2013; Donchin, 1981). If the probe P300s are larger than irrelevant P300s, this is used to infer criminal guilt. Many studies have found highly accurate detection of memories with this technique, and variations of the CIT are used in real life criminal investigations (Matsuda et al., 2019) and sold commercially (Farwell, 2012). However, recent research has exposed vulnerabilities of this method to factors that impede memory detection (Bergström et al., 2013; Hu et al., 2015; Meijer et al., 2016; Rosenfeld et al., 2004). Thus, P300-based CIT may not be as accurate as originally thought, which may have dire real-life consequences.

One important issue that has not yet been investigated is the effect of healthy aging on P300-based forensic memory detection, despite evidence that aging has strong effects on memory (Cabeza et al., 2018; Nyberg et al., 2012) and ERPs associated with recognition (Friedman, 2013). In this latter line of research, ERPs are typically measured while participants complete an episodic old/new recognition test where they need to discriminate previously encountered (old) from new stimuli. In these tests, old and new items are typically presented just once and in equal proportions, and thus neither stimulus category is subjectively rare. In such designs, younger adults typically show an enlarged parietal ERP positivity around 0.5-1s during recognition of old compared to new stimuli, referred to as the parietal old–new effect (Rugg & Curran, 2007; Wilding, 2000). In contrast, older adults often show reduced (Ally et al., 2008; Duarte et al., 2006; Wang et al., 2012) or even reversed (i.e. negative, Horne et al., 2020; Li et al., 2004) parietal old–new effects, often despite intact behavioural test accuracy. This literature thus suggests that recognition-related parietal ERPs change with aging, which could potentially affect P300-based memory detection.

The relationship between parietal old–new effects and P300 recognition effects is unclear, however. Although they have similar time-course and morphology, there are many differences between the paradigms used to elicit these ERP effects, and they are typically interpreted as functionally distinct. Parietal old–new effects have been argued to reflect conscious recollection of contextual details from an event (Vilberg & Rugg, 2009; Wilding, 2000), whereas P300 effects in oddball tasks are typically interpreted as related to attention, working memory and/or decision making (Donchin, 1981; Polich, 2007). However, other evidence suggest both effects are related to attentional orienting or decision making (e.g. Dywan et al., 1998; Yang et al., 2019; although see Herron et al., 2003). Consistent with a functional link, some findings suggest that oddball P300 effects are also reduced in older age (Polich et al., 1985; van Dinteren et al., 2014), but in those studies the P300 response was not based on recognition of information from a specific event such as in the CIT. Therefore, based on previous literature, it is not known whether CIT P300 memory detection is affected by healthy aging. In the current study, we recorded ERPs from older and younger participants while they completed both a standard episodic old/new recognition test and a CIT to investigate if P300-based CIT is less accurate in older suspects, and if so, how this reduced accuracy relates to changes in parietal old–new ERP effects during episodic recognition.

We also complemented the above ERP analyses by applying time-frequency decomposition to the same EEG data, to measure oscillations associated with recognition. This technique has several advantages over ERPs that may be especially relevant for understanding aging-related memory changes. First, it can be used to measure induced EEG oscillations that are not phase-locked to an event (e.g. Tallon-Baudry & Bertrand, 1999) and are therefore less sensitive to neural timing variability across trials (“temporal jitter”), which is increased with aging (Tran et al., 2016) and can reduce the amplitude of ERP components. Time-frequency analysis is therefore particularly useful for estimating high frequency EEG oscillations that are less likely to be phase-locked across trials than slow frequency EEG responses. Second, because this technique separates the EEG into non-overlapping frequency bands, it may be better able to dissociate different cognitive processes than ERPs, which reflect a summation of event-related EEG across all frequency bands and is therefore more susceptible to component overlap. Such component overlap has been suggested as a potential contributor to age-differences in episodic recognition-related ERPs (Allen et al., 2020; Dulas & Duarte, 2013) and it is therefore important to investigate if recognition-related EEG oscillations are also affected by aging.

In young adults, episodic recognition is typically related to increased power (i.e. synchronization) in the theta frequency band (~4-7Hz), which often peaks across the parietal scalp around the same time as parietal ERP positivities occur (i.e. ~0.5-1s after stimulus onset; for reviews see Herweg et al., 2020; Hsieh & Ranganath, 2014; Nyhus & Curran, 2010). In addition, episodic recognition is often also associated with reduced power (i.e. desynchronization) in the alpha (~8-12Hz) and beta (~13-30Hz) frequency bands, with different topography depending on the content of retrieval and type of retrieval processing (see Hanslmayr et al., 2016). Less is known regarding how aging affects the oscillatory correlates of recognition, but recent studies have shown relatively similar patterns of retrieval-related theta synchronization and alpha desynchronization for both younger and older adults during episodic recognition (Allen et al., 2020; Karlsson et al., 2020; Strunk et al., 2017), although some of these studies also found age differences in alpha desynchronization. EEG oscillations are not typically analysed in practical applications of the CIT, but other types of “oddball” research has shown that orienting responses/P300 ERP effects are also associated with theta synchronization (Bachman & Bernat, 2018; Harper et al., 2017; Keller et al., 2017) and at least one prior study has shown alpha desynchronization to be associated with P300s (Bernat et al., 2007). The effects of aging on EEG in oddball tasks have not been studied to the same extent, but one study found that both theta synchronization and alpha/beta desynchronization was enhanced in older compared to younger adults (Ho et al., 2012).

In the current study, we investigated for the first time if P300-based CIT is less accurate in older suspects, and if so, how this reduced accuracy relates to changes in parietal old–new ERP effects during episodic recognition. Are these two effects both reduced in our older compared to younger group? Do the same individuals show reductions in both effects, as would be expected if they share underlying mechanisms? We also applied time-frequency analysis to our EEG data to assess whether aging-related reductions in parietal ERP positivities were associated with changes to EEG oscillations in theta, alpha and beta frequency bands. Based on prior literature we expected to observe similar theta synchronization and alpha/beta desynchronization old–new effects in both the recognition memory test and the CIT in the younger group. If recognition-related ERP differences between younger and older adults were due to increased temporal jitter and/or component overlap, then the older group was expected to show EEG oscillation effects that were similar to those in the younger group. In contrast, if reduced parietal ERP positivities in the older group were due to a reduced engagement of recognition-related neurocognitive processes, then the older group would be expected to show reduced oscillatory markers of recognition to parallel the ERP results. Answering these questions thus have important implications for forensic applications, and for theoretical accounts of the role of different brain processes in recognition memory.

## Methods

### Participants

Participants were recruited from a database of research volunteers or adverts placed around the University of Kent, and were compensated at approx. £7.50/hour. The study was approved by the Research Ethics Committee of the School of Psychology at the University of Kent. The final sample consisted of 30 young adults (*M*_age_ = 22, *SD*_age_ = 4, range = 18-33, 19 females) and 30 old adults (*M*_age_ = 69, *SD*_age_ = 6, range = 60-80, 22 females). This sample size was chosen to have 0.85 power to detect a *d*=0.8 effect size, or 0.8 power to detect a *d*=0.75 effect size (we were primarily interested in detecting large effects of aging on ERPs and behaviour, since only large effects have practical implications for forensic memory detection). An additional four younger adults and one older adult were excluded due to poor EEG recording quality. All participants were right-handed, native English speakers, had no psychiatric or neurological diagnosis, and reported not taking any medicine that affects cognition. All participants completed the Montreal Cognitive Assessment (MOCA; Nasreddine et al., 2005) after the experimental tasks. Operation Span (OSPAN; Unsworth et al., 2005) and Wechsler abbreviated scale of intelligence (WASI-II; Wechsler, 2011) scores were either gathered from existing datasets or participants attended a second session to complete these tests. Older adults had higher age-corrected WASI IQ scores than younger adults, whereas the younger adults performed higher in the OSPAN working memory test (Table 1). The Bayes Factors indicated moderate support for no difference between age groups in the MOCA.

**Table 1.** Cognitive test results

|  | The Montreal<br>cognitive assessment<br>(MOCA) | The Wechsler abbreviated<br>Scale of intelligence II<br>(WASI) | OSPAN Partial Score |
| --- | --- | --- | --- |
| Younger Group | 28.03<br>[27.31, 28.76] | 109.63<br>[105.66, 113.61] | 67.03<br>[64.05, 70.01] |
| Older Group | 27.70<br>[26.96, 28.44] | 116.37<br>[112.37, 120.36] | 54.47<br>[47.98, 60.95] |
| *t*-value | -.66 | 2.44 | -3.60 |
| *p*-value | .51 | .018 | < .001 |
| Cohen's *d* | .17 | .63 | .93 |
| Bayes Factors<br>(BF<sub>01</sub>) | 3.31/1 | 1/3.03 | 1/44.13 |

Note: values show mean scores and 95% confidence interval (in square brackets) for each cognitive test. Group differences were assessed with independent t-tests and Bayes Factors for the null hypothesis over the experimental hypothesis of a group difference in either direction (BF<sub>01</sub>).

### Materials, Design and Procedure

All participants first carried out a mock crime, which was followed by a brief computerised word reading test to ensure that they were able to read and understand words at the pace required during the CIT. Next, participants completed an incidental word encoding task, which was followed by the CIT, and finally a recognition test for the words from the encoding Task.

#### Mock Crime

Participants were given a map and instructed to go to a kitchen and steal an object (a ring) inside a cupboard. The word “ring” was never used in the instructions to ensure that participants acquired the crime-related memory through enacting the mock crime. Participants were asked to examine the stolen object and bring it to the EEG lab, but were not told that their memory for the object would later be tested. Next, they were asked if they had noticed anything else inside the cupboard (a watch, which was to be used as the target in the CIT). If they had not, they were asked to go back to the kitchen and look again.

#### Encoding Task

Participants then carried out an incidental encoding task, in which 64 words were displayed on a computer screen in a randomised order for 3000ms each, preceded by a 600ms fixation cross. Participants were not told that their memory for the words would later be tested, and instead rated the words for “pleasantness” using button presses on a response box. The 64 words were drawn from a larger pool of 128 words from the MRC Psycholinguistic database (Wilson, 1988), of which the remaining 64 words would be used as foils in a subsequent surprise recognition test (counterbalanced across participants). Strong semantic associations between words within the recognition task and also with words in the CIT were avoided.

#### Concealed Information Test

The CIT involved repeatedly presenting participants with six words on a computer screen: a crime-relevant probe (“ring”), a target (“watch”) and four irrelevants (“phone”, “keys”, “necklace”, “wallet”). Participants were informed that an object had been stolen from a nearby kitchen and they needed to complete a test to detect if they were guilty of the theft. It was explained that words would be shown on the screen and they should press buttons on a response box to indicate whether they recognised objects from the kitchen, while their brain activity was recorded. They were told that they should avoid responding “yes” to the word corresponding to the stolen object (i.e. the probe “ring”) in order to avoid being found guilty, and only respond “yes” when they saw the target word “watch”. Each of the six words were presented 60 times each in a randomised order. Trials began with a black screen for a jittered duration between 500-1000ms, followed by a 500ms fixation cross before the word was displayed for 300ms in white on a black background. The word was followed by a blank screen for 1000ms and button presses were accepted for 1750ms from word onset. Six attention checks were pseudo-randomly interspersed every 42-78 CIT trials, which required participants to verbally respond with the most recently presented word. There was no significant difference in number of failed attention checks between the younger and older groups (younger: *M* = .23, 95% CI = [.07, .39], range = 0-1; older: *M* = .47, 95% CI = [.18,.76], range = 0-3; *t*(58) = 1.44, *p* = .155, BF_01_ = 1.60/1).

#### Recognition Test

The CIT was followed by a recognition test, where the 64 “old” words from the incidental encoding task were shown on the computer screen randomly intermixed with 64 “new” foil words and participants indicated whether or not they recognised the words using the response box. Words were displayed for 3000ms and preceded by a 600ms fixation cross.

#### Post Experiment Questionnaire

After the computerised tasks, participants responded to a few questions assessing their self-reported experiences of the mock crime and CIT. This questionnaire was also used to verify that all participants knew what object they had stolen and had seen the probe word (i.e. “ring”) during the CIT (which they all had).

### EEG Recording and Pre-processing

EEG was recorded from 64 scalp electrodes (plus vertical and horizontal EOG) positioned in accordance with the extended 10-20 system using an actiCAP system (Brain Products Gmbh) at 500Hz with a 0.05-70Hz bandwidth, and with impedances kept below 25 kΩ. The EEG was pre-processed offline using the EEGLAB toolbox (Delorme & Makeig, 2004) in MATLAB, with a standard pipeline (similar to Bergström et al., 2013) involving filtering with a 0.3 Hz-30Hz bandpass, re-referencing to the average of both mastoids, and removing non-brain artefacts with Independent Component Analysis and manual screening. Epochs were extracted from –200ms to 1700ms post stimulus presentation in both the recognition test and the CIT, and baseline corrected using the 200ms pre-stimulus interval. ERPs were computed by averaging the raw EEG epochs for each condition and participant separately, including all trials independent of accuracy. Mean trial numbers across ERP conditions ranged between 55-223, all participants contributed at least 44 trials, and mean trial numbers were highly similar across the two age groups (mean differences ranged between 0-2 trials).

#### Time-Frequency Decomposition

The same single-trial EEG data that were averaged to form ERPs were also submitted to time-frequency decomposition using a complex Morlet wavelet transform as implemented in FieldTrip (Oostenveld et al., 2011). The pre-stimulus interval in each epoch (600ms in the recognition memory test and 1000ms in the CIT) was mirrored and prepended to the original pre-stimulus data to provide a long enough interval for reliably estimating the pre-stimulus baseline power of lower frequencies (*cf.* Vogelsang et al., 2018). The decomposition was performed between 4-30 Hz in frequency steps of 1 Hz, time steps of 5ms and a wavelet width of three cycles. The epochs were truncated to –825 to 1300ms in both the recognition memory test and the CIT to avoid edge artefacts and make the analysis time-window comparable across tests. Oscillatory power in each frequency band was normalised to a dB scale using a baseline period between –825 to –375ms pre-stimulus. This baseline period was selected to avoid artefactual “bleeding” of post-stimulus EEG activity into the baseline because of the low temporal resolution of low frequency wavelets (as in Allen et al., 2020; Vogelsang et al., 2018).

### Statistical Analysis Approach

For key behavioural and focal ERP analysis, we used frequentist *t*-tests. Cohen’s *d* effect sizes were calculated as the difference between means divided by the pooled standard deviation (Dunlap, Cortina, Vaslow, & Burke, 1996). *T*-tests were supplemented with Bayes Factors in order to assess the relative evidence for the alternative versus the null hypothesis. The Bayes Factor (BF_01_) is a ratio that contrasts the likelihood that the data would occur under the null (H_0,_ no difference between groups/conditions/no difference from 0) versus alternative (H_1_, a group difference/condition difference/a difference from 0) hypotheses. Ratios close to 1 are only considered weakly/anecdotally supportive of one hypothesis over the other, whereas BF_01_ >3 are typically interpreted as substantial evidence in support of H_0_ over H_1_, and BF_01_ <1/3 are interpreted as substantial evidence in support of H_1_ over H_0_. All Bayes factors were calculated with two-tailed tests, using JASP (JASP Team, 2017) with recommended default priors (a Cauchy distribution with centre = 0, r = 0.707).

#### Focal Analysis of Individual ERP Peaks

We first conducted a focal analysis of the electrodes where the P300 and parietal old–new effects are typically maximal (mid-parietal (Pz) and left parietal (P3) sites respectively). In order to control for potential individual and age differences in the timing of recognition-related ERP effects, we followed previous CIT research (e.g. Hu et al., 2015) and applied a sliding window to individual ERP conditions to measure parietal positivities as the mean amplitude of the 100ms window with the most positive amplitude between 300-1000ms. The amplitudes of recognition-related parietal positivities were then computed by subtracting the control conditions from the recognition conditions (i.e. old–new, probe–irrelevant, target– irrelevant). These recognition-related ERP effects were then compared between groups with independent *t*-tests to test for age differences. ERP effects were compared to 0 with one-sample *t*-tests to assess their presence within each age group.

#### Global Analysis of ERP Data

Analysing ERP data from only a few electrode sites and time-windows may overlook effects occurring at other scalp locations and time points, which may be problematic when comparing age groups since the topography and timing of recognition-related ERP effects can change with age (Friedman, 2013). We therefore complemented the focal analysis with global data-driven analyses where we included all scalp electrodes and all time points between 0-1700ms, and controlled for multiple comparisons using nonparametric cluster-based permutation tests with the FieldTrip toolbox (Oostenveld et al., 2011) in MATLAB. This analysis involved first performing *t*-tests at every ERP data sample to estimate significant differences (uncorrected at *α*=.05) between conditions/groups. Significant samples that were adjacent in time and space (spanning at least two electrodes) were grouped into clusters, and their *t*-values were summed into one cluster-level *t-*value. The false positive rate for the full spatiotemporal data matrix was controlled by testing the observed cluster-level *t*-values against a null distribution of cluster-level *t*-statistics created by permutation resampling, where ERP data were randomly allocated to conditions for all participants and clusters were recalculated for each resample (5000 in total). *P*-values were calculated as the proportion of the randomization null distribution exceeding the observed maximum cluster-level test statistic (i.e. the Monte-Carlo *p*-value). This analysis enabled us to identify significant clusters extending over time and electrodes (see Maris & Oostenveld, 2007). We tested if there was a group difference in each ERP effect of interest (old–new, probe– irrelevant and target–irrelevant), and also assessed presence of those ERP effects within each age group.

#### Global Analysis of Time-Frequency Data

Similarly to the global ERP analysis, the time-frequency data were analysed with nonparametric cluster-based permutation tests (Maris & Oostenveld, 2007). All post-stimulus time points (0-1300ms), electrodes and frequencies (4-30 Hz) were included in the analyses. Clusters were formed between significant neighbouring samples in the time, frequency and electrode dimensions. The same parameters were used as in the global analysis of the ERP data. We first tested group differences in the relevant contrasts (old–new, probe–irrelevant and target–irrelevant). Next, we conducted within group cluster-based permutation tests to investigate the presence of EEG oscillation effects within each age group.

Additional analyses assessed whether age groups differed in terms of individual-level ERP-based memory detection, and trial-by-trial temporal jitter of parietal ERP peaks. The individual level results mirrored the ERP findings reported in the next section, and the temporal jitter analysis showed no significant differences in temporal variability of parietal peaks between age groups (these analyses will be available on OSF when this preprint is published in a journal).

## Results

### Behavioural Results

Recognition test and CIT performance is shown in Table 2. The younger group had faster reaction times than the older group on the recognition test, but there were no significant age differences in recognition test discrimination or response bias. The older group were more accurate than the younger group for probes on the CIT, but there were no other significant age differences in CIT performance.

**Table 2.** Behavioural performance on the Recognition Test and the CIT.

| Group | Recognition Test |  |  |  | Concealed Information Test |  |  |  |  |  |
| --- | --- | --- | --- | --- | --- | --- | --- | --- | --- | --- |
|  | Discrimination and Response Bias |  | Reaction Times |  | Accuracy |  |  | Reaction Times |  |  |
|  | *Pr* | *Br* | Old | New | Target | Probe | Irrelevant | Target | Probe | Irrelevant |
| Younger Group | .83<br>[.79, .87] | .39<br>[.31, .48] | 879<br>[831, 928] | 939<br>[879, 1000] | .91<br>[.88, .94] | .97<br>[.96, .99] | .99<br>[.99, 1.00] | 548<br>[504, 591] | 475<br>[430, 521] | 462<br>[425, 499] |
| Older Group | .78<br>[.73, .83] | .41<br>[.32, .50] | 1030<br>[965, 1095] | 1017<br>[975, 1059] | .94<br>[.91, .96] | .99<br>[.99, 1.00] | .99<br>[.99, 1.00] | 566<br>[532, 600] | 501<br>[460, 541] | 493<br>[466, 521] |
| *t*-value | -1.57 | .28 | -3.81 | -2.17 | -1.63 | -2.03 | -.34 | -.67 | -.86 | -1.40 |
| *p*-value | (.12) | (.78) | (<.001) | (.034) | (.11) | (.047) | (.74) | (.51) | (.40) | (.17) |
| Cohen's *d* | .40 | .07 | .98 | .56 | .42 | .56 | .09 | .17 | .22 | .36 |
| Bayes Factors<br>(BF<sub>01</sub>) | 1.37/1 | 3.69/1 | 1/76.92 | 1/1.81 | 1.26/1 | 1/1.45 | 3.63/1 | 3.16/1 | 2.80/1 | 1.69/1 |

Note: Values show means and 95% confidence intervals (in square brackets). Group differences were assessed with independent *t*-tests and Bayes Factors for the null hypothesis over the experimental hypothesis (BF<sub>01</sub>). The *Pr* measure provides a single measure of participants' ability to correctly discriminate between old and new items ( $Pr = \text{hits} - \text{false alarms}$ ). The *Br* measure provides an estimate of participants' bias toward responding "old" vs. "new", independently of accuracy ( $Br = \text{false alarms} / (1 - Pr)$ ; see Snodgrass & Corwin, 1988).

Self-reports (Table 3) showed that crime memories were equally likely to come to mind automatically for both age groups, but the younger group were more motivated than the older group to “beat” the CIT and try to appear innocent.

**Table 3.** CIT Questionnaire results

|  | Question |  |  |  |  |
| --- | --- | --- | --- | --- | --- |
|  | Nervousness during the mock crime (0-6) | Extent to which crime memories came to mind during the CIT (0-6) | Motivation to beat the CIT test (0-6) | Used strategies to beat the CIT (1= yes, 0 = no) | (If answered yes to previous question) Confidence in strategies (0-6) |
| Younger Group | 2.00<br>[1.52, 2.48] | 3.62<br>[2.94, 4.31] | 4.37<br>[3.88, 4.85] | .66<br>[.47, .84] | 3.00<br>[2.32, 3.68] |
| Older Group | 1.57<br>[1.11, 2.02] | 3.33<br>[2.62, 4.05] | 3.10<br>[2.38, 3.82] | .40<br>[.10, .70] | 3.22<br>[2.15, 4.29] |
| *t*-value | -1.34 | -.59 | -2.98 | -1.46 | .39 |
| *p*-value | .19 | .56 | .004 | .15 | .70 |
| Cohen's *d* | .35 | .15 | .77 | .38 | .16 |
| Bayes Factors (BF<sub>01</sub>) | 1.81/1 | 3.27/1 | 1/9.35 | 1/2.74 | 2.55/1 |

Note: Participants answered yes/no questions or rated their mock crime/CIT experiences on a 0-6 scale, with higher numbers indicating more of the measured experience than lower numbers. Values show means and 95% confidence interval (in square brackets) of self-report ratings from the post experimental questionnaire. Group differences were assessed with independent t-tests and Bayes Factors for the null hypothesis over the experimental hypothesis of a group difference in either direction (BF<sub>01</sub>).

### Concealed Information Test ERPs

First, we analysed the CIT ERPs to test if recognition-related P300s were modulated by aging.

#### Focal analysis of CIT ERPs

The ERP waveforms from the analysed electrode (Pz) and individual target–irrelevant P300 and probe–irrelevant P300 differences are illustrated in Fig. 1. Consistent with predictions, the probe–irrelevant P300 effect was significantly larger in the younger group compared to older group, *t*(58) = 2.79, *p* = .007, *d* = .72, BF_01_ = 1/6.12. In fact, while the effect was highly significant in the younger group, *t*(29) = 3.03, *p* = .005, *d* = .25, BF_01_ = 1/8.05, it was not significant in the older group, and the Bayes Factor indicated moderate support for an absence of a P300 probe–irrelevant effect in the older group, *t*(29) = −.27, *p* = .79, *d* = .018, BF_01_ = 4.97/1.

**Fig. 1.**
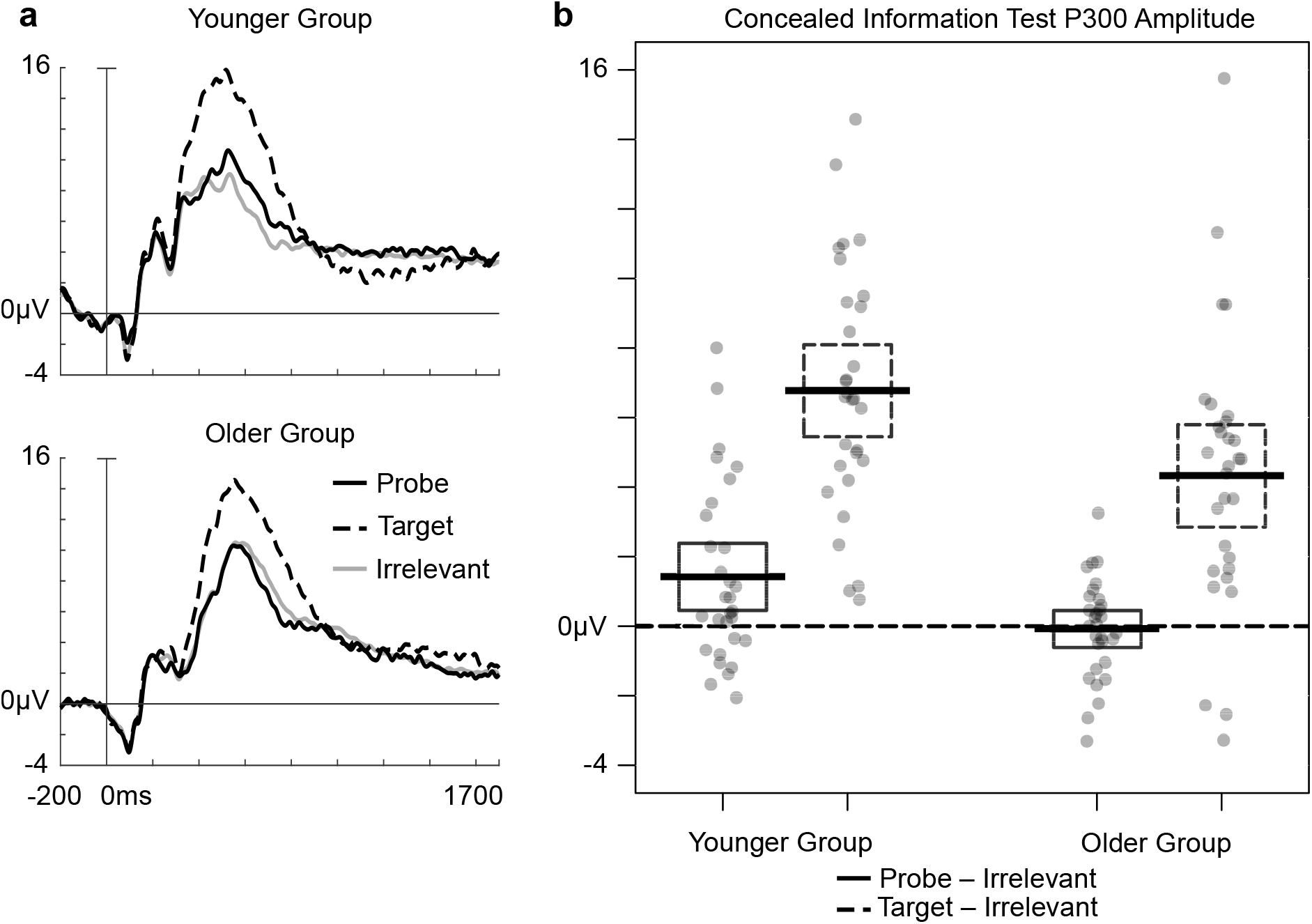
Focal analysis of CIT ERPs. **a.** Grand average ERP waveforms from midline parietal electrode site (Pz) for target, probe and irrelevant items in the CIT. **b.** Individually estimated amplitudes of the probe-irrelevant and target-irrelevant P300 effects at electrode Pz. The scatter dots show individual P300 effect amplitudes. The thick lines show the group means and the boxes depict the 95% confidence interval of the group means.

The target–irrelevant P300 effect was also significantly larger in the younger group compared to the older group, *t*(58) = 2.53, *p* = .014, *d* = .65, BF_01_ = 1/3.58, although importantly it was highly significant in both groups (younger group: *t*(29) = 10.53, *p* < .001, *d* = 1.13, BF_01_ = 1/9.63e+12; older group: *t*(29) = 5.99, *p* < .001, *d* = .93, BF_01_ = 1/1.13e+4). Thus, it was not the case that older adults showed a complete absence of P300 effects, since they did show a typical enlarged target P300 response. Target P300s and probe P300s were however reduced by a similar extent in the older compared to younger group, as the target– probe P300 difference did not significantly interact with age, *t*(58) = −.92, *p* = .36, *d* = .24, BF_01_ = 2.67/1.

#### Global analysis of CIT ERPs

The results from the global analysis of the CIT ERPs are shown in Fig. 2. As predicted and consistent with the focal analysis, the probe–irrelevant P300 effect was significantly larger in the younger group compared with the older group (*p* = .002). This effect corresponded to a mid-posterior cluster between approximately 510-870ms. Follow up analyses confirmed that there was a typical probe–irrelevant P300 effect in the younger group (*p* = .007) as indicated by a positive mid-posterior cluster between approximately 450-840ms. In contrast, the probe– irrelevant P300 effect was not only absent in the older group, but even reversed, as shown by significantly more negative ERPs for probes than irrelevants (*p* = .013) between approximately 620-780ms with a broad scalp distribution. This analysis thus revealed a qualitatively different probe effect in the older group, compared to the typical young probe P300.

**Fig. 2.**
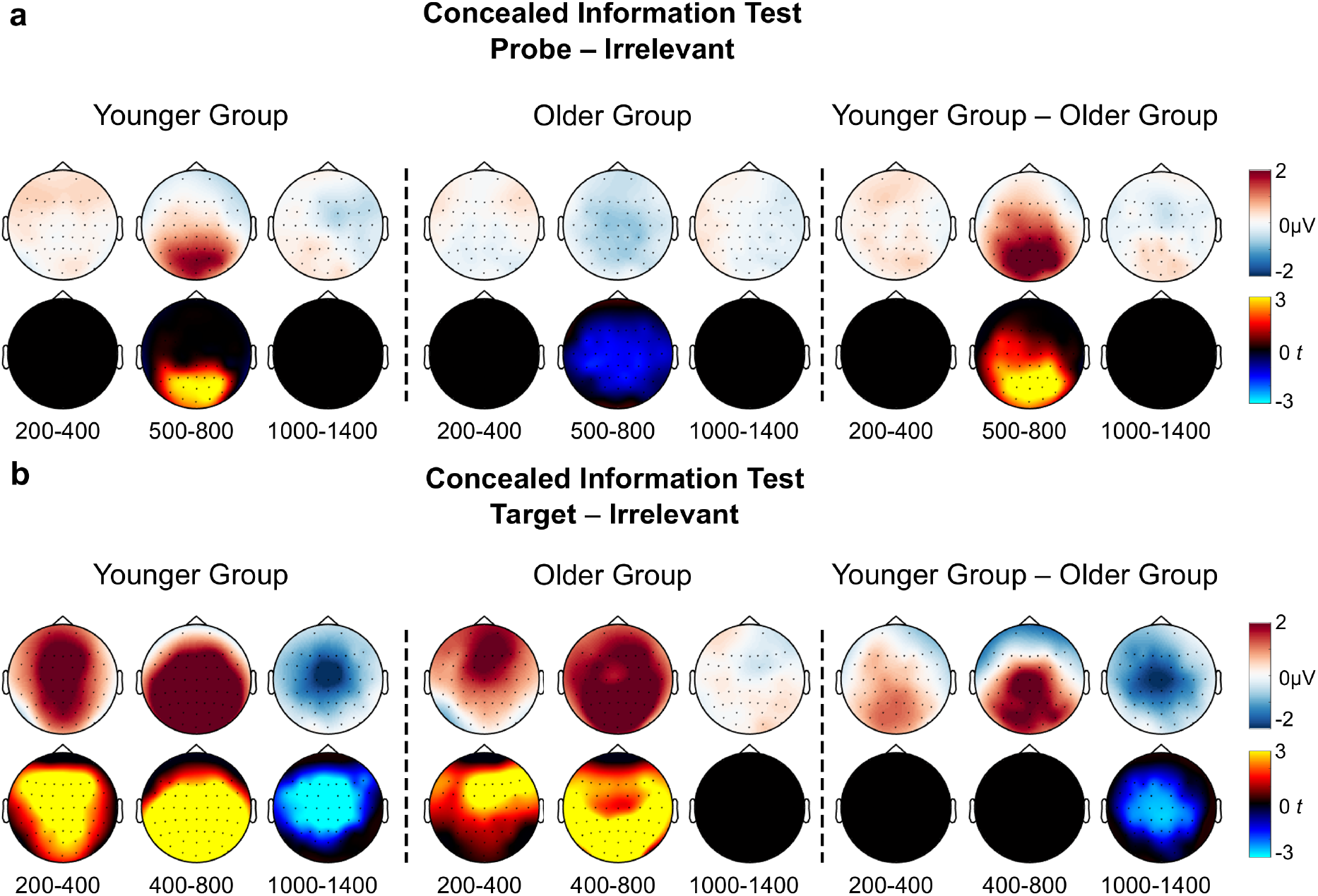
Global analysis of CIT ERPs. Topographical maps of amplitude differences (top rows, blue/white/red colourmap) and *t*-values for the differences (bottom rows, cold/black/hot colour map) between **a**) target and irrelevant conditions and **b**) probe and irrelevant conditions in the CIT. Only significant clusters (*p* < .025 at the cluster level, which corresponds to a two-tailed α*=*.05) are shown in the t-statistic topographical maps.

The cluster-based permutation test indicated that the target–irrelevant P300 effect was only trend-level larger in the younger group compared to the older group (*p* = .032, meaning it just failed to exceed the *α*-level). This cluster was maximal across central and parietal sites and lasted between approximately 560-690ms. The target–irrelevant late negativity was highly significantly larger in the younger group compared to the older group (*p* < .001). This effect corresponded to a central negative cluster between approximately 1100-1470ms. Follow-up analyses indicated both a positive broadly distributed target–irrelevant P300 (*p* < .001) between around 200-900ms, and a centrally distributed target–irrelevant late negativity (*p* = .001) between approximately 970-1470ms in the younger group. A target–irrelevant P300 positivity was also seen in the older group (*p* < .001) between 220-950ms, but this group did not show a significant target–irrelevant late negativity. Thus, the target P300 was more similar across age groups, appearing to only numerically differ in a quantitative way.

### Concealed Information Test EEG Oscillations

Next, we analysed EEG oscillations from the CIT to test if recognition-related theta, alpha and beta oscillations were modulated by aging.

#### Probe–Irrelevant EEG Oscillations

As can be seen in Fig. 3, there was a larger alpha/beta (around 8-30 Hz) desynchronization effect for probes than irrelevant items in the younger group compared to the older group between approximately 450-860ms (*p* = .023). The corresponding negative cluster effect had a posterior distribution. This age group difference was driven by a widespread probe– irrelevant alpha/beta desynchronization effect within the young group (peaking around 10-30 Hz, between approximately 480-1000ms, *p* < .001), but there was no similar effect in the older group. In the younger group, there was also a widespread probe–irrelevant theta (~4-8Hz) synchronization effect around 60-1050ms (*p* = .02) that was not present within the older group, but the interaction with age group for this effect did not survive the cluster-based permutation test correction for false positives^1^. Thus, paralleling the ERP analysis, the EEG oscillation results revealed significantly reduced and even absent probe-irrelevant differences in the older group compared to the younger group.

**Fig. 3.**
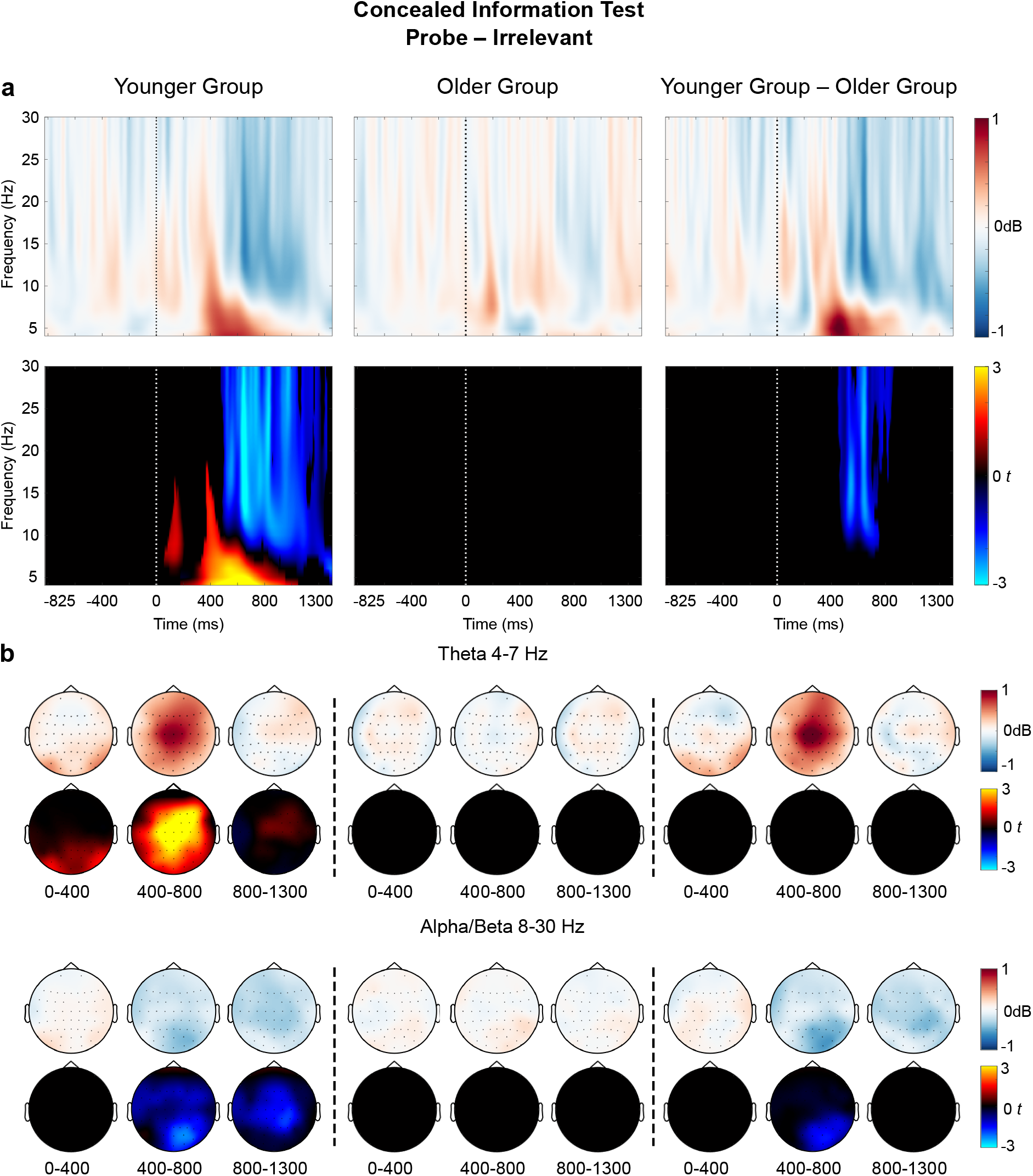
Results of a global time-frequency analysis of probe–irrelevant differences in the Concealed Information Test. **a**. Time-Frequency plots of probe–irrelevant differences in power (top row, blue/white/red colourmap) and t-values for the differences (bottom row, cold/black/hot colour map). The time-frequency plots show averaged data over all electrodes. Only significant clusters (*p* < .025 at the cluster level) are shown in the t-statistic time-frequency plots, and note that the pre-stimulus interval was not analysed statistically. **b.** Topographic maps of probe–irrelevant differences in the Theta (4-7Hz, top two rows) and Alpha/Beta (8-30Hz, bottom two rows) frequency bands. Within each frequency band, the top row (blue/white/red colourmap) illustrates the topographical distributions of power differences. The bottom row within each frequency band (cold/black/hot colour map) shows t-values for the differences. Only significant clusters (*p* < .025 at the cluster level) are shown in the t-statistic topographic maps and there is consequently only a partial correspondence between the power and the t-value plots.

#### Target – Irrelevant EEG Oscillations

The results from the time-frequency analysis of the target–irrelevant contrast in the CIT are depicted in Fig. 4. The cluster analysis indicated that there was significantly higher power for the target–irrelevant difference in the younger group compared to the older group between around 40-1300ms (peaking around 600-1000ms), in a cluster that spanned across all frequencies (*p* < .001). This positive cluster was due to an early larger theta synchronization effect in the younger group (around 40-400ms) followed by enhanced broadband desynchronization in the older group (approximately 400-1300ms). Within the younger group, there was an early widespread target–irrelevant theta synchronization effect (maximal between 4-10Hz) between 0-1070ms, peaking around 200-550ms (*p* = .007). This effect was followed by a posterior broadband desynchronization effect between approximately 410-1300ms, corresponding to a negative cluster between 5-30Hz (*p* < .001). The older group had a similar broadband desynchronization effect between approximately 300-1300ms (*p* < .001), that had a broadly distributed topography and included all analysed frequencies (4-30Hz). Although a theta synchronization effect was numerically visible in the older group between around 400-800ms across the fronto-central scalp, this did not survive the cluster-based permutation test correction for false positives.^2^ Thus, consistent with the ERP analysis there were robust oscillatory target-irrelevant differences in both age groups indicating that the reduction in EEG effects in the older group are specific to the memory related probe-irrelevant contrast.

**Fig. 4.**
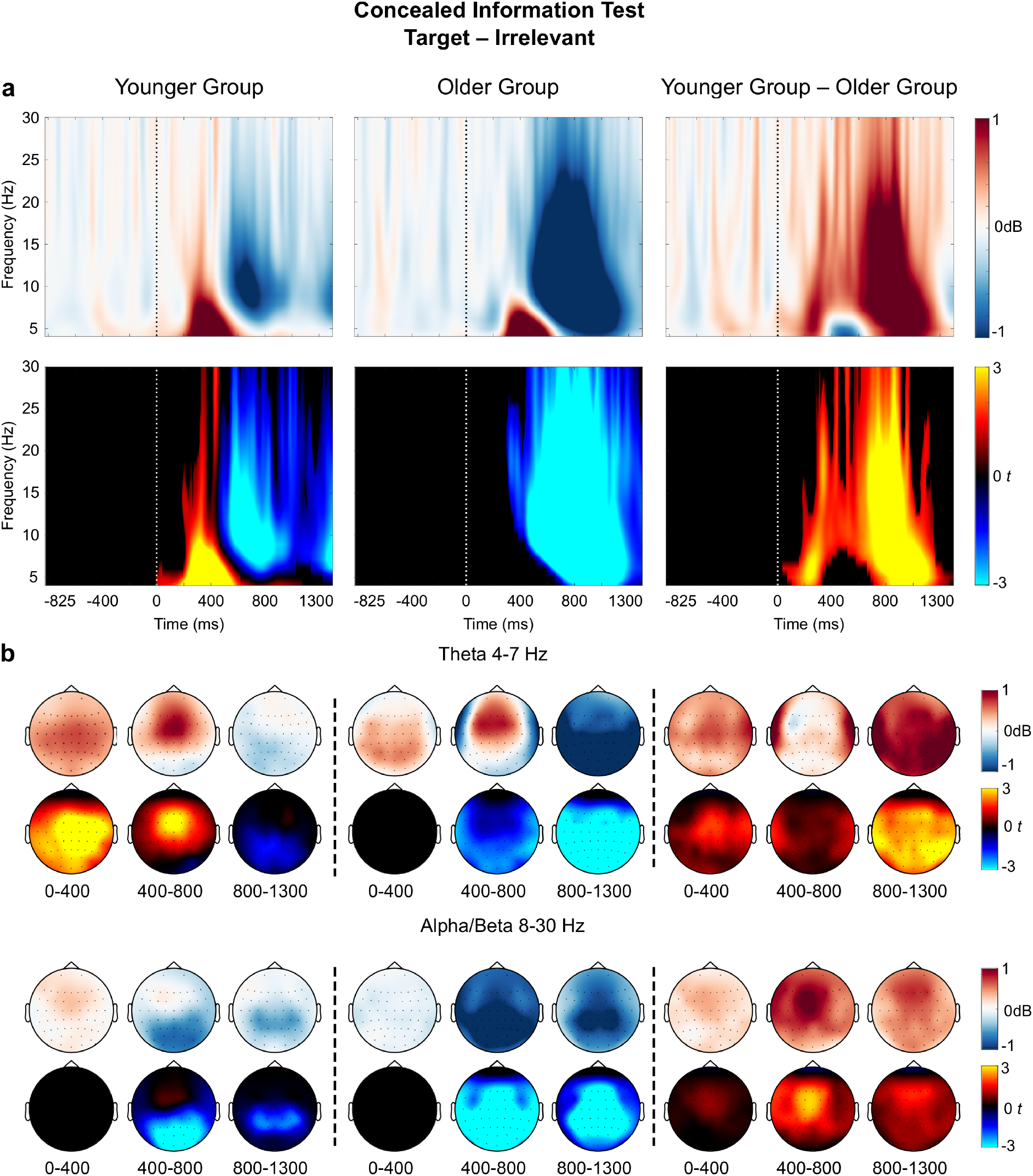
Results of a global time-frequency analysis of target–irrelevant differences in the Concealed Information Test. **a**. Time-Frequency plots of target–irrelevant differences in power (top row, blue/white/red colourmap) and t-values for the differences (bottom row, cold/black/hot colour map). The time-frequency plots show averaged data over all electrodes. Only significant clusters (*p* < .025 at the cluster level) are shown in the t-statistic time-frequency plots, and note that the pre-stimulus interval was not analysed statistically. **b.** Topographic maps of target–irrelevant differences in the theta (4-7Hz, top two rows) and alpha/beta (8-30Hz, bottom two rows) frequency bands. Within each frequency band, the top row (blue/white/red colourmap) illustrates the topographical distributions of power differences. The bottom row within each frequency band (cold/black/hot colour map) shows t-values for the differences. Only significant clusters (*p* < .025 at the cluster level) are shown in the t-statistic topographic maps and there is consequently only a partial correspondence between the power and the t-value plots.

### Recognition Test ERPs

In the next step, we analysed the parietal old–new ERP effects on the recognition test to determine if these also were affected by aging.

#### Focal Analysis of Recognition Test ERPs

The grand average ERPs from the analysed electrode (P3) and the individually estimated peak amplitudes of left parietal old–new effects are depicted in Fig. 5.

**Fig. 5.**
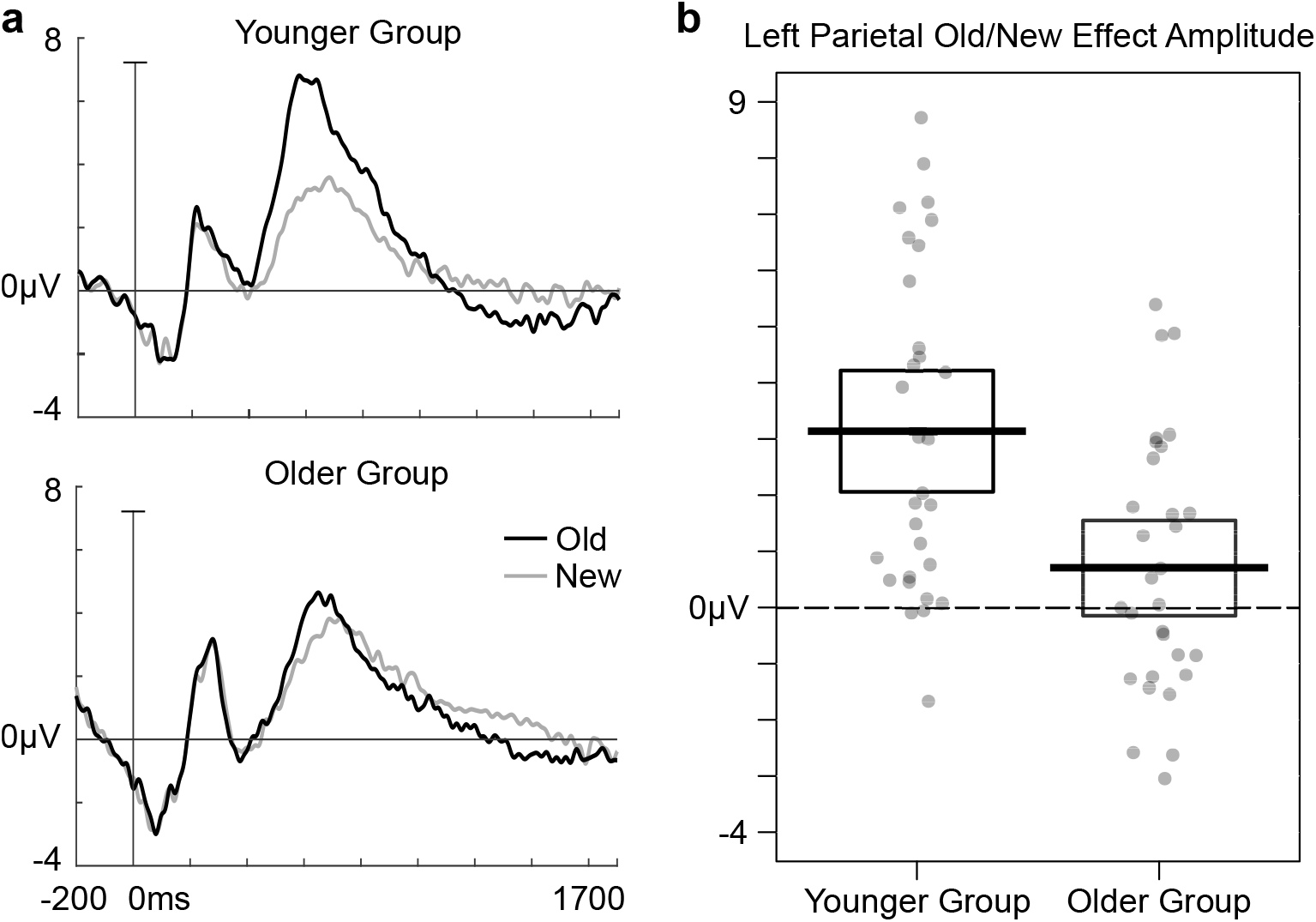
Focal analysis of recognition test ERPs. **a.** Grand average ERPs from a left parietal electrode site (P3) for old and new items in the recognition memory test. **b.** Individually estimated amplitudes of the left parietal old–new effect at electrode P3. Dots show individual participant’s amplitudes and are randomly scattered across the x-axis for visualisation purposes. The thick lines show the group means and the boxes depict the 95% confidence interval of the group means.

As predicted, the left parietal old–new effect was significantly larger in the younger group than in the older group, *t*(58) = 3.62, *p* < .001, *d* = .93, BF_01_ = 1/45.46. The effect was significant within the younger group, *t*(29) = 5.94, *p* < .001, *d* = .79, BF_01_ 1/9960, but not in the older group, *t*(29) = 1.70, *p* = .10, *d* = .21, BF_01_ = 1.43/1.

#### Global Analysis of Recognition Test ERPs

The results from the global analysis of the recognition test ERPs are illustrated in Fig. 6. Mirroring the results from the focal analysis, the cluster-based permutation test also showed that the left parietal old–new effect was significantly larger in the younger group than the older group (*p* = .020). This left posterior cluster extended from approximately 450-660ms post-stimulus presentation.

**Fig. 6.**
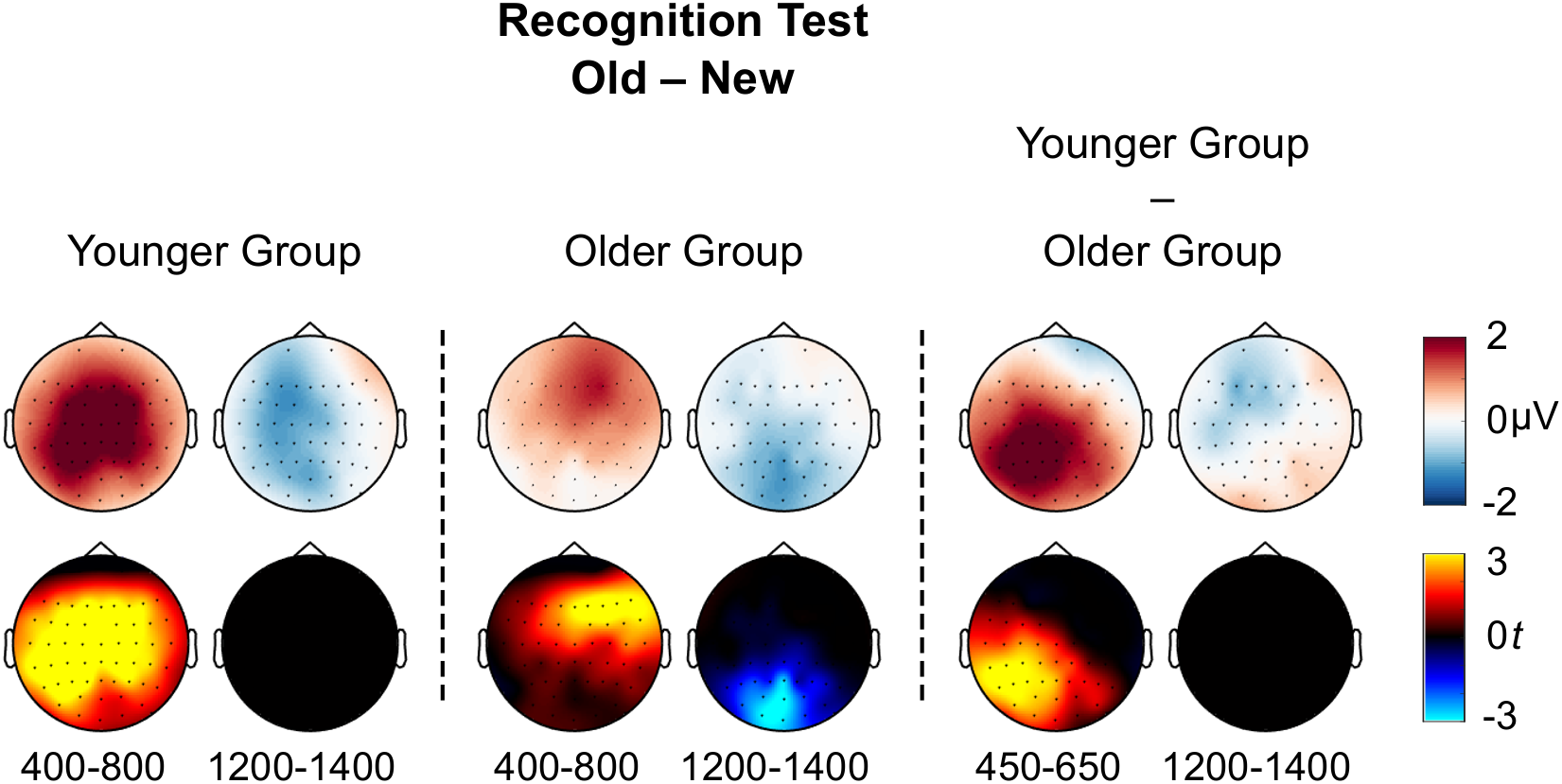
Global analysis of recognition test ERPs. Topographic maps of old–new amplitude differences (top rows, blue/white/red colourmap) and *t*-values for the differences (bottom rows, cold/black/hot colourmap) in the recognition memory test. Only significant clusters (*p* < .025 at the cluster level, which corresponds to a two-tailed α*=*.05) are shown in the t-statistic topographic maps.

Follow up analysis showed that there was a significant positive going old–new effect in the younger group (*p* < .001) that was maximal across parietal and central sites between approximately 420-910ms. There was also a trend towards a late negative old–new effect (*p* = .033), with a broad scalp distribution maximal between approximately 1160-1410ms (but note that this effect is not shown in the t-value plot). Interestingly, there was also a positive going old–new effect in the older group (*p* = .003), but this effect had a right anterior, rather than left posterior distribution between approximately 420-780ms. In addition, the older group also had a significant late negative old–new effect (*p* = .024) corresponding to a mid-posterior cluster between approximately 1220-1450ms.

### Recognition Test EEG Oscillations

We next analysed EEG oscillations from the recognition test to assess whether these were modulated by ageing in a parallel way to recognition test ERPs (Fig. 7).

**Fig. 7.**
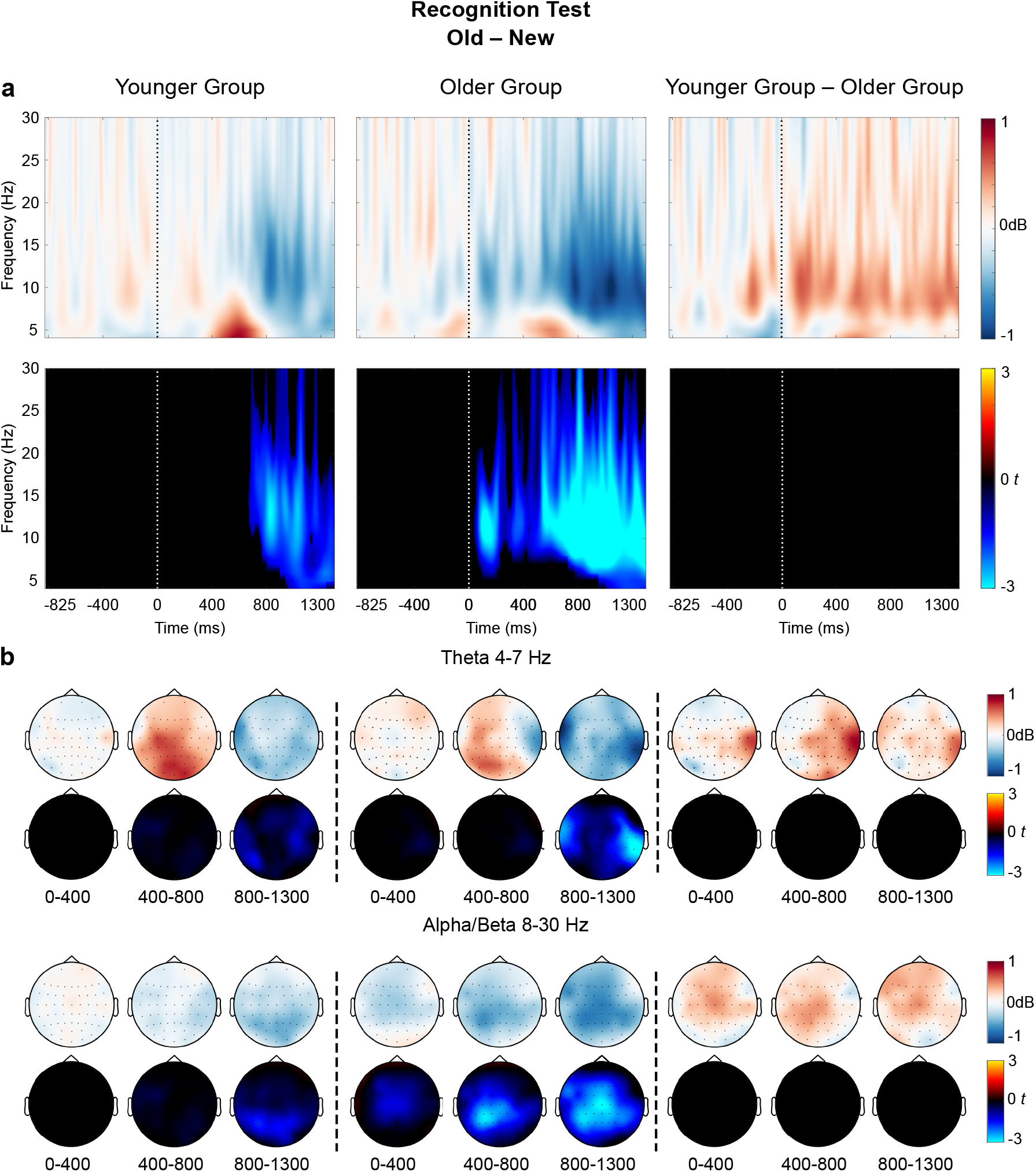
Results of a global time-frequency analysis of old–new differences in the Recognition Test EEG data. **a**. Time-Frequency plots of old–new differences in power (top row, blue/white/red colourmap) and t-values for the differences (bottom row, cold/black/hot colour map). The time-frequency plots show averaged data over all scalp electrodes. Only significant clusters (*p* < .025 at the cluster level) are shown in the t-statistic time-frequency plots; note that pre-stimulus interval was not analysed statistically. **b.** Topographic maps of old–new differences in the Theta (4-7Hz, top two rows) and Alpha/Beta (8-30Hz, bottom two rows) frequency bands. Within each frequency band, the top row (blue/white/red colourmap) illustrates the topographical distributions of power differences. The bottom row within each frequency band (cold/black/hot colour map) shows t-values for the differences. Only significant clusters (*p* < .025 at the cluster level) are shown in the t-statistic topographic maps and there is consequently only a partial correspondence between the power and the t-value plots.

In contrast to the ERP results, the cluster-based permutation tests indicated that there were no significant groups differences in oscillatory old–new effects in the recognition memory test. In the younger group, there was a broadly distributed desynchronization old–new effect in frequencies between around 5-20 Hz, that was maximal between 670-1300ms (*p* = .005). There was a similar broadly distributed desynchronization old–new effect in frequencies around 7-20Hz in the older group between approximately 40-1300ms, peaking around 550-1300ms (*p* < .001). The predicted recognition-related theta synchronization effects that are numerically visible in Fig. 7 between ~400-800ms post stimulus across the parietal scalp did not survive the cluster-based permutation test correction for false positives, which is primarily sensitive to effects that are broadly distributed across time, space and frequency domains^3^.

### Relationship between the left parietal old–new effect and the probe–irrelevant P300 effect

In a final analysis, we investigated if individual differences in the size of recognition-related parietal positivities correlated across the recognition test and CIT. In the younger group, there was a positive correlation between the size of the left parietal old–new effect and the probe– irrelevant P300 (Kendall’s tau B = .26, *p* = .049, BF_01_ = 1/1.55). In contrast, in the older group the Bayes Factors indicated moderate support for no correlation between the left parietal old–new effect and the probe–irrelevant P300 effect (Kendall’s tau B = .-.10, *p* = .44, BF_01_ = 3.12/1).

## Discussion

We investigated the effects of healthy aging on parietal P300s in an ERP-based forensic memory detection test, and compared these with parietal old–new effects during an episodic recognition test. Consistent with predictions, older adults showed reduced recognition-related parietal ERP positivities on both tests compared to younger adults, despite both groups showing highly accurate and similar behavioural performance. Reduced recognition-related parietal ERP positivities in older age have been previously found in episodic memory tests (Ally et al., 2008; Duarte et al., 2006; Friedman, 2013; Horne et al., 2020; Li et al., 2004; Wang et al., 2012), but have not been shown before in P300-based detection of concealed information, which typically focuses on young (~18-20 years old) student populations (e.g. Bergström et al., 2013; Bowman et al., 2013; Hu et al., 2015; Meijer et al., 2014; Rosenfeld, 2020). These novel findings raise concerns about the validity of ERP-based forensic memory detection with older adults, and questions about the functional role of parietal ERP positivities in episodic recognition.

Aging-related reductions in P300s elicited by mock crime probes relative to irrelevant control stimuli were found in both a focal analysis of parietal ERPs with individually adjusted time-windows, and in a comprehensive data-driven analysis across all time-points and scalp locations. These analyses showed typical probe–irrelevant P300 effects in the younger group but no evidence of probe–irrelevant P300 effects in the older group, whereas both age groups showed highly significant, enlarged P300s to target stimuli compared to irrelevant control stimuli. In the CIT framework (e.g. Meijer et al., 2014; Rosenfeld, 2020), probe P300s are thought to index automatic recognition of incriminating information. At face value, it might therefore appear that these healthy older adults without memory impairments did not recognise the word “ring” as crime-relevant despite having “stolen” a real ring less than 30 minutes earlier, a task that is thought to create a sensorimotor rich, vivid autobiographical memory in young adults (e.g. Hu et al., 2015). However, this conclusion is contradicted by participants’ self-reports, since both age groups reported recognising the word “ring” while undertaking the CIT and that this had automatically brought memories of the mock crime to mind. Furthermore, the global analysis revealed a reversed parietal ERP pattern in the older group, with enhanced negative ERPs for probes compared to irrelevant stimuli, showing that probes did elicit a different brain response than irrelevants in the older group, but this response was qualitatively different from the brain response in the younger group.

EEG oscillations during the CIT mirrored the P300 ERP pattern, with reduced recognition-related oscillatory effects in the probe-irrelevant contrast in the older group compared to the younger group, despite highly significant target–irrelevant EEG oscillation effects in both groups. These target–irrelevant effects were largely consistent with prior literature on the oddball paradigm, with early theta synchronization followed by alpha/beta desynchronization for targets compared to irrelevant items (Bachman & Bernat, 2018; Bernat et al., 2007; Harper et al., 2017; Keller et al., 2017). Interestingly, this latter alpha/beta desynchronization effect for targets was enhanced in older adults, showing that the absence of oscillatory effects in the probe–irrelevant contrast in the same group cannot simply be explained by noisier EEG measures in older than younger participants, or a failure of older participants to attend to the task.

Similar to the Probe-Irrelevant P300, Parietal ERP positivities for previously seen words during the old/new recognition test were also reduced in the older group, despite older participants demonstrably recognising those words as shown by equally high behavioural task accuracy across age groups. These results also indicate that an absence of parietal ERP positivities cannot be used as evidence for an absence of recognition in older adults. A global analysis revealed that the older group did have a positive old–new effect within the typical ~0.5-1s time-window, but this positivity had a right anterior topography (*cf.* Horne et al., 2020). Older adults’ episodic word recognition could thus be detected with ERPs, but it involved a different pattern of brain activity compared to the younger group, who showed a typical left parietal peak.

In contrast, the time-frequency results showed that older and younger adults had relatively similar EEG oscillations on the episodic recognition test, despite large ERP differences. This pattern replicates our recent study on aging effects on recognition memory, where we also found a significantly reduced left parietal old–new effect in the ERPs in an older group compared to a younger group, but no age differences in oscillatory old–new effects in theta or alpha bands (Allen et al., 2020). The results from the episodic recognition test therefore suggest that oscillation analysis of EEG can reveal recognition-related brain activity that is obscured in older people’s ERPs.

Why were recognition-related parietal ERP positivities reduced in older compared to younger adults? First, several explanations can be ruled out. It was not simply the case that EEG measurements were noisy overall in older adults, since they still showed a number of other highly significant ERP and oscillation effects. In addition, aging-related parietal ERP reductions were found even when adjusting for individual differences in peak timing, and supplementary analyses showed no evidence of group differences in trial-by-trial temporal jitter of parietal peaks, meaning age differences cannot be explained by delayed or more variable parietal peak timing in the older group (*cf.* Bielak et al., 2014; Murray et al., 2019; van Dinteren et al., 2014b). Instead, the results suggest that older adults did have intact ability to recognise old stimuli and crime probes, but that the brain processes they engaged during recognition differed from the young group. Interestingly, while the size of parietal peaks on the CIT and old/new recognition test correlated across individuals within the younger group, they did not correlate across individuals within the older group. This pattern suggests that parietal ERP positivities may be functionally related across CIT and episodic recognition tests (cf. Dywan et al., 1998; Yang et al., 2019), but that aging-related reductions to those positivities may have been driven by different causes on the two tests.

Reduced or reversed recognition-related parietal ERP effects in older age may be caused by “component overlap” with parietally distributed ERP negativities that summate and “cancel out” positive peaks at the scalp (e.g. Dulas & Duarte, 2013; Horne et al., 2020). Such late posterior negativities (LPNs) are often enhanced in situations that require response monitoring or careful evaluation of memory details (Mecklinger et al., 2016). In older adults, LPNs may index processes that are recruited to improve performance in cognitively demanding situations in order to compensate for aging-related memory impairments (Friedman, 2013). In our study, reduced parietal old–new ERP effects in older age may have been primarily caused by LPN effects related to compensatory processes. Since such LPN effects are manifest as slow EEG drifts, they are more likely to influence ERP measures than measures of relatively fast EEG oscillations. Consistent with this account, the older and younger adults showed similar recognition-related oscillation effects, suggesting that the different age groups also recruited some overlapping episodic recognition processes.

In contrast to the findings on the episodic recognition test, aging-related P300 modulations on the CIT may have been influenced also by a different mechanism. When subjectively rare CIT probes are recognised based on memory of a recent event, this likely involves both episodic retrieval (Rugg & Curran, 2007) and orienting-related processes (Polich, 2007), that may both contribute to P300 amplitudes through summation at the scalp (Bergström et al., 2013; Herron et al., 2003). The orienting response is sensitive to arousal and motivational factors and is thought to be mediated by noradrenaline and dopamine systems (Nieuwenhuis et al., 2011). Such neurotransmitter functioning is altered in older age and linked with aging-related memory changes (Mather, 2016), which might have contributed to probe P300 reductions in our older group. This suggestion is consistent with the finding that probes did not elicit a significant difference in EEG oscillations compared to irrelevant items within the older group, suggesting that the older group was simply not showing a strong orienting response to probes, in contrast to the younger adults. Interestingly, younger adults reported higher motivation to “beat” the CIT and appear innocent than the older adults, consistent with prior findings in young adults that motivation to appear innocent can paradoxically enhance orienting responses (Meijer et al., 2014). Future research should test how arousal and motivation affects recognition-related ERPs and EEG oscillations across ages, in order to better understand the neural mechanisms underlying EEG and ERP markers of memory and how they are influenced by aging.

In conclusion, here we show for the first time that EEG-based forensic memory detection is impaired for older adults, and that aging-related EEG changes on a concealed information test are accompanied by aging-related changes to episodic recognition-related ERPs, but that these changes may be driven by non-overlapping mechanisms. We also demonstrate that EEG-based forensic memory detection in older age is impaired even with a comprehensive analysis approach with multiple complementary measures that compensate for potential age differences in frequency, timing and topography of brain responses. Our findings raise concerns regarding the validity of EEG-based forensic memory detection tests, and suggest that their practical application (e.g. Farwell, 2012; Matsuda et al., 2019) is premature.

## Author Contributions

Z.M. Bergström, R. Hellerstedt, A. Moccia and H. Bowman developed the research questions and designed the experiments. The data was collected by A. Moccia and C.M. Brunskill. Data were analysed and interpreted by R. Hellerstedt and Z.M. Bergström. R. Hellerstedt and Z.M. Bergström drafted the manuscript and all co-authors provided critical revisions. All authors approved the final version of the manuscript for submission.

## Open Practices Statement

The experiment was not formally preregistered. Data and materials will be shared on OSF when this preprint is published in a journal.

## Footnotes

1 If we analyse the Theta band oscillations separately from Alpha/Beta bands, then the Theta synchronization effect does show a significant interaction with age group (*p* < .001), and is significant larger in the Younger than Older group.

2 For the record, if we analyse the Theta band oscillations separately from Alpha/Beta bands, then the Theta synchronization target-irrelevant effect is significant also within the Older group (*p* < .001), although it still interacts with age group (there were two significant positive clusters in the theta band, one between 60-350ms, *p* = .018 and one between 670-1230ms, p< .001).

3 If we analyse the Theta band oscillations separately from Alpha/Beta bands, then the Theta synchronization effect is significant only within the younger group (*p* = .006), but does not show a significant interaction with age group.

## Notes

### Competing Interest Statement

The authors have declared no competing interest.

## References

Allen, J., Hellerstedt, R., Sharma, D., & Bergström, Z. M. (2020). Distraction by Unintentional Recognition: Neurocognitive Mechanisms and Effects of Aging. Psychology and Aging, 35(5), 639–653. doi: 10.1037/pag0000398

Ally, B. A., Simons, J. S., McKeever, J. D., Peers, P. V., & Budson, A. E. (2008). Parietal contributions to recollection: Electrophysiological evidence from aging and patients with parietal lesions. Neuropsychologia, 46(7), 1800–1812. https://doi.org/10.1016/j.neuropsychologia.2008.02.026

Bachman, M. D., & Bernat, E. M. (2018). Independent contributions of theta and delta time-frequency activity to the visual oddball P3b. International Journal of Psychophysiology: Official Journal of the International Organization of Psychophysiology, 128, 70–80. https://doi.org/10.1016/j.ijpsycho.2018.03.010

Bergström, Z. M., Anderson, M. C., Buda, M., Simons, J. S., & Richardson-Klavehn, A. (2013). Intentional retrieval suppression can conceal guilty knowledge in ERP memory detection tests. Biological Psychology, 94(1), 1–11. https://doi.org/10.1016/j.biopsycho.2013.04.012

Bernat, E. M., Malone, S. M., Williams, W. J., Patrick, C. J., & Iacono, W. G. (2007). Decomposing delta, theta, and alpha time–frequency ERP activity from a visual oddball task using PCA. International Journal of Psychophysiology: Official Journal of the International Organization of Psychophysiology, 64(1), 62–74. https://doi.org/10.1016/j.ijpsycho.2006.07.015

Bielak, A. A. M., Cherbuin, N., Bunce, D., & Anstey, K. J. (2014). Intraindividual variability is a fundamental phenomenon of aging: Evidence from an 8-year longitudinal study across young, middle, and older adulthood. Developmental Psychology, 50(1), 143–151. https://doi.org/10.1037/a0032650

Bowman, H., Filetti, M., Janssen, D., Su, L., Alsufyani, A., & Wyble, B. (2013). Subliminal salience search illustrated: EEG identity and deception detection on the fringe of awareness. PloS One, 8(1), e54258. https://doi.org/10.1371/journal.pone.0054258

Cabeza, R., Albert, M., Belleville, S., Craik, F. I. M., Duarte, A., Grady, C. L., Lindenberger, U., Nyberg, L., Park, D. C., Reuter-Lorenz, P. A., Rugg, M. D., Steffener, J., & Rajah, M. N. (2018). Maintenance, reserve and compensation: The cognitive neuroscience of healthy ageing. Nature Reviews Neuroscience, 19(11), 701–710. https://doi.org/10.1038/s41583-018-0068-2

Delorme, A., & Makeig, S. (2004). EEGLAB: An open source toolbox for analysis of single-trial EEG dynamics including independent component analysis. Journal of Neuroscience Methods, 134(1), 9–21. https://doi.org/10.1016/j.jneumeth.2003.10.009

Donchin, E. (1981). Surprise!… Surprise? Psychophysiology, 18(5), 493–513. https://doi.org/10.1111/j.1469-8986.1981.tb01815.x

Duarte, A., Ranganath, C., Trujillo, C., & Knight, R. T. (2006). Intact Recollection Memory in High-performing Older Adults: ERP and Behavioral Evidence. Journal of Cognitive Neuroscience, 18(1), 33–47. https://doi.org/10.1162/089892906775249988

Dulas, M. R., & Duarte, A. (2013). The influence of directed attention at encoding on source memory retrieval in the young and old: An ERP study. Brain Research, 1500, 55–71. https://doi.org/10.1016/j.brainres.2013.01.018

Dywan, J., Segalowitz, S. J., & Webster, L. (1998). Source Monitoring: ERP Evidence for Greater Reactivity to Nontarget Information in Older Adults. Brain and Cognition, 36(3), 390–430. https://doi.org/10.1006/brcg.1997.0979

Farwell, L. A. (2012). Brain fingerprinting: A comprehensive tutorial review of detection of concealed information with event-related brain potentials. Cognitive Neurodynamics, 6(2), 115–154. https://doi.org/10.1007/s11571-012-9192-2

Friedman, D. (2013). The Cognitive Aging of Episodic Memory: A View Based on the Event-Related Brain Potential. Frontiers in Behavioral Neuroscience, 7. https://doi.org/10.3389/fnbeh.2013.00111

Hanslmayr, S., Staresina, B. P., & Bowman, H. (2016). Oscillations and Episodic Memory: Addressing the Synchronization/Desynchronization Conundrum. Trends in Neurosciences, 39(1), 16–25. https://doi.org/10.1016/j.tins.2015.11.004

Harper, J., Malone, S. M., & Iacono, W. G. (2017). Theta- and delta-band EEG network dynamics during a novelty oddball task. Psychophysiology, 54(11), 1590–1605. https://doi.org/10.1111/psyp.12906

Herron, J. E., Quayle, A. H., & Rugg, M. D. (2003). Probability effects on event-related potential correlates of recognition memory. Cognitive Brain Research, 16(1), 66–73. https://doi.org/10.1016/S0926-6410(02)00220-3

Herweg, N. A., Solomon, E. A., & Kahana, M. J. (2020). Theta Oscillations in Human Memory. Trends in Cognitive Sciences, 24(3), 208–227. https://doi.org/10.1016/j.tics.2019.12.006

Ho, M.-C., Chou, C.-Y., Huang, C.-F., Lin, Y.-T., Shih, C.-S., Han, S.-Y., Shen, M.-H., Chen, T.-C., Liang, C., Lu, M.-C., & Liu, C.-J. (2012). Age-related changes of task-specific brain activity in normal aging. Neuroscience Letters, 507(1), 78–83. https://doi.org/10.1016/j.neulet.2011.11.057

Horne, E. D., Koen, J. D., Hauck, N., & Rugg, M. D. (2020). Age differences in the neural correlates of the specificity of recollection: An event-related potential study. Neuropsychologia, 140, 107394. https://doi.org/10.1016/j.neuropsychologia.2020.107394

Hsieh, L.-T., & Ranganath, C. (2014). Frontal midline theta oscillations during working memory maintenance and episodic encoding and retrieval. NeuroImage, 85 Pt 2, 721–729. https://doi.org/10.1016/j.neuroimage.2013.08.003

Hu, X., Bergström, Z. M., Bodenhausen, G. V., & Rosenfeld, J. P. (2015). Suppressing Unwanted Autobiographical Memories Reduces Their Automatic Influences: Evidence From Electrophysiology and an Implicit Autobiographical Memory Test. Psychological Science, 26(7), 1098–1106. https://doi.org/10.1177/0956797615575734

Karlsson, A. E., Wehrspaun, C. C., & Sander, M. C. (2020). Item recognition and lure discrimination in younger and older adults are supported by alpha/beta desynchronization. BioRxiv, 2020.04.22.055764. https://doi.org/10.1101/2020.04.22.055764

Keller, A. S., Payne, L., & Sekuler, R. (2017). Characterizing the roles of alpha and theta oscillations in multisensory attention. Neuropsychologia, 99, 48–63. https://doi.org/10.1016/j.neuropsychologia.2017.02.021

Li, J., Morcom, A. M., & Rugg, M. D. (2004). The effects of age on the neural correlates of successful episodic retrieval: An ERP study. Cognitive, Affective, & Behavioral Neuroscience, 4(3), 279–293. https://doi.org/10.3758/CABN.4.3.279

Lykken, D. T. (1959). The GSR in the detection of guilt. Journal of Applied Psychology, 43(6), 385–388. https://doi.org/10.1037/h0046060

Maris, E., & Oostenveld, R. (2007). Nonparametric statistical testing of EEG- and MEG-data. Journal of Neuroscience Methods, 164(1), 177–190. https://doi.org/10.1016/j.jneumeth.2007.03.024

Mather, M. (2016). The Affective Neuroscience of Aging. Annual Review of Psychology, 67(1), 213–238. https://doi.org/10.1146/annurev-psych-122414-033540

Matsuda, I., Ogawa, T., & Tsuneoka, M. (2019). Broadening the Use of the Concealed Information Test in the Field. Frontiers in Psychiatry, 10, 24. https://doi.org/10.3389/fpsyt.2019.00024

Mecklinger, A., Rosburg, T., & Johansson, M. (2016). Reconstructing the past: The late posterior negativity (LPN) in episodic memory studies. Neuroscience and Biobehavioral Reviews, 68, 621–638. https://doi.org/10.1016/j.neubiorev.2016.06.024

Meijer, E. H., klein Selle, N., Elber, L., & Ben-Shakhar, G. (2014). Memory detection with the Concealed Information Test: A meta analysis of skin conductance, respiration, heart rate, and P300 data. Psychophysiology, 51(9), 879–904. https://doi.org/10.1111/psyp.12239

Meijer, E. H., Verschuere, B., Gamer, M., Merckelbach, H., & Ben-Shakhar, G. (2016). Deception detection with behavioral, autonomic, and neural measures: Conceptual and methodological considerations that warrant modesty. Psychophysiology, 53(5), 593–604. https://doi.org/10.1111/psyp.12609

Morcom, A. M. (2016). Mind Over Memory: Cuing the Aging Brain. Current Directions in Psychological Science, 25(3), 143–150. https://doi.org/10.1177/0963721416645536

Murray, J. G., Ouyang, G., & Donaldson, D. I. (2019). Compensation of Trial-to-Trial Latency Jitter Reveals the Parietal Retrieval Success Effect to be Both Variable and Thresholded in Older Adults. Frontiers in Aging Neuroscience, 11. https://doi.org/10.3389/fnagi.2019.00179

Nasreddine, Z. S., Phillips, N. A., Bédirian, V., Charbonneau, S., Whitehead, V., Collin, I., Cummings, J. L., & Chertkow, H. (2005). The Montreal Cognitive Assessment, MoCA: A brief screening tool for mild cognitive impairment. Journal of the American Geriatrics Society, 53(4), 695–699. https://doi.org/10.1111/j.1532-5415.2005.53221.x

Nieuwenhuis, S., de Geus, E. J., & Aston-Jones, G. (2011). The anatomical and functional relationship between the P3 and autonomic components of the orienting response. Psychophysiology, 48(2), 162–175. https://doi.org/10.1111/j.1469-8986.2010.01057.x

Nyberg, L., Lövdén, M., Riklund, K., Lindenberger, U., & Bäckman, L. (2012). Memory aging and brain maintenance. Trends in Cognitive Sciences, 16(5), 292–305. https://doi.org/10.1016/j.tics.2012.04.005

Nyhus, E., & Curran, T. (2010). Functional Role of Gamma and Theta Oscillations in Episodic Memory. Neuroscience and Biobehavioral Reviews, 34(7), 1023–1035. https://doi.org/10.1016/j.neubiorev.2009.12.014

Oostenveld, R., Fries, P., Maris, E., & Schoffelen, J.-M. (2011). FieldTrip: Open source software for advanced analysis of MEG, EEG, and invasive electrophysiological data. Computational Intelligence and Neuroscience, 2011, 156869. https://doi.org/10.1155/2011/156869

Polich, J. (2007). Updating P300: An integrative theory of P3a and P3b. Clinical Neurophysiology, 118(10), 2128–2148. https://doi.org/10.1016/j.clinph.2007.04.019

Polich, J., Howard, L., & Starr, A. (1985). Effects of age on the P300 component of the event-related potential from auditory stimuli: Peak definition, variation, and measurement. Journal of Gerontology, 40(6), 721–726. https://doi.org/10.1093/geronj/40.6.721

Rosenfeld, J. P. (2020). P300 in detecting concealed information and deception: A review. Psychophysiology, 57(7), e13362. https://doi.org/10.1111/psyp.13362

Rosenfeld, J. P., Soskins, M., Bosh, G., & Ryan, A. (2004). Simple, effective countermeasures to P300-based tests of detection of concealed information. Psychophysiology, 41(2), 205–219. https://doi.org/10.1111/j.1469-8986.2004.00158.x

Rugg, M. D., & Curran, T. (2007). Event-related potentials and recognition memory. Trends in Cognitive Sciences, 11(6), 251–257. https://doi.org/10.1016/j.tics.2007.04.004

Snodgrass, J. G., & Corwin, J. (1988). Pragmatics of measuring recognition memory: Applications to dementia and amnesia. Journal of Experimental Psychology: General, 117(1), 34–50. https://doi.org/10.1037/0096-3445.117.1.34

Strunk, J., James, T., Arndt, J., & Duarte, A. (2017). Age-related changes in neural oscillations supporting context memory retrieval. Cortex; a Journal Devoted to the Study of the Nervous System and Behavior, 91, 40–55. https://doi.org/10.1016/j.cortex.2017.01.020

Tallon-Baudry, C., & Bertrand, O. (1999). Oscillatory gamma activity in humans and its role in object representation. Trends in Cognitive Sciences, 3(4), 151–162. https://doi.org/10.1016/S1364-6613(99)01299-1

Tran, T. T., Hoffner, N. C., LaHue, S. C., Tseng, L., & Voytek, B. (2016). Alpha phase dynamics predict age-related visual working memory decline. NeuroImage, 143, 196–203. https://doi.org/10.1016/j.neuroimage.2016.08.052

Unsworth, N., Heitz, R. P., Schrock, J. C., & Engle, R. W. (2005). An automated version of the operation span task. Behavior Research Methods, 37(3), 498–505. https://doi.org/10.3758/bf03192720

van Dinteren, R., Arns, M., Jongsma, M. L. A., & Kessels, R. P. C. (2014). P300 development across the lifespan: A systematic review and meta-analysis. PloS One, 9(2), e87347. https://doi.org/10.1371/journal.pone.0087347

Vilberg, K. L., & Rugg, M. D. (2009). Functional Significance of Retrieval-related Activity in Lateral parietal cortex: Evidence from fMRI and ERPs. Human Brain Mapping, 30(5), 1490–1501. https://doi.org/10.1002/hbm.20618

Vogelsang, D. A., Gruber, M., Bergström, Z. M., Ranganath, C., & Simons, J. S. (2018). Alpha oscillations during incidental encoding predict subsequent memory for new “foil” information. Journal of Cognitive Neuroscience, 30(5), 667–679.

Wang, T. H., de Chastelaine, M., Minton, B., & Rugg, M. D. (2012). Effects of age on the neural correlates of familiarity as indexed by ERPs. Journal of Cognitive Neuroscience, 24(5), 1055–1068. https://doi.org/10.1162/jocn_a_00129

Wechsler, D. (2011). Wechsler Abbreviated Scale of Intelligence–Second Edition (WASI-II). NCS Pearson.

Wilding, E. L. (2000). In what way does the parietal ERP old/new effect index recollection? International Journal of Psychophysiology: Official Journal of the International Organization of Psychophysiology, 35(1), 81–87. https://doi.org/10.1016/s0167-8760(99)00095-1

Wilson, M. (1988). MRC psycholinguistic database: Machine-usable dictionary, version 2.00. Behavior Research Methods, Instruments, & Computers, 20(1), 6–10. https://doi.org/10.3758/BF03202594

Yang, H., Laforge, G., Stojanoski, B., Nichols, E. S., McRae, K., & Köhler, S. (2019). Late positive complex in event-related potentials tracks memory signals when they are decision relevant. Scientific Reports, 9(1), 9469. https://doi.org/10.1038/s41598-019-45880-y

